# Identifying key residues in intrinsically disordered regions of proteins using machine learning

**DOI:** 10.1101/2022.12.09.519711

**Authors:** Wen-Lin Ho, Hsuan-Cheng Huang, Jie-rong Huang

## Abstract

Conserved residues in protein homolog sequence alignments are structurally or functionally important. For intrinsically disordered proteins (IDPs) or proteins with intrinsically disordered regions (IDRs), however, alignment often fails because they lack a steric structure to constrain evolution. Although sequences vary, the physicochemical features of IDRs may be preserved in maintaining function. Therefore, a method to retrieve common IDR features may help identify functionally important residues. We applied un-supervised contrastive learning to train a model with self-attention neuronal networks on human IDR orthologs. During training, parameters were optimized to match sequences in ortholog pairs but not in other IDRs. The trained model successfully identifies previously reported critical residues from experimental studies, especially those with an overall pattern (e.g. multiple aromatic residues or charged blocks) rather than short motifs. This predictive model can therefore be used to identify potentially important residues in other proteins.

**Availability and implementation:** The training scripts are available on GitHub (https://github.com/allmwh/IFF). The training datasets have been deposited in an Open Science Framework repository (https://osf.io/jk29b). The trained model can be run from the Jupyter Notebook in the GitHub repository using Binder (mybinder.org). The only required input is the primary sequence.

## Introduction

DNA/RNA sequences and the proteins they encode carry their evolutionary history, and multiple sequence alignment methods can reveal phylogenetic relationships. For instance, our extinct Neanderthal and Denisovan cousins were identified from the DNA extracted in ancient bones [1, 2]; prokaryotic ribosomal 16S RNA sequences contributed to the discovery of Archaea domain [3]; the tracing of myoglobin and hemoglobin protein sequences back to their globin origin is another textbook example [4, 5]. Protein structures also provide insights into how proteins have evolved, being conserved in some cases despite changes in the primary sequence. For example, the structural similarity between the motor domains of kinesin and myosin hints that they have a common ancestor, despite low sequence identity [6]. The shape of a protein also constrains how it evolves and functionally important residues are conserved. Indeed, when sequence conservation levels are mapped onto 3D structures, the most conserved residues are typically found in key locations, such as the folding core [7] or catalytic sites [8].

However, these structural constraints on evolution do not apply to intrinsically disordered proteins (IDPs) or proteins with intrinsically disordered regions (IDRs), and as a result, the sequences of these proteins, which represent more than half of the proteome [9], vary more widely than do those of their folded counterparts (see example in Supplementary Figure S1). Although some structural evolutionary restraints still apply to some IDRs, especially those that undergo folding-upon-binding [10, 11], the evolution of IDRs is mainly constrained by function. One recently recognized function of IDRs is their ability to undergo liquid-liquid phase separation (LLPS) [12, 13]. This mechanism contributes to the formation of membraneless organelles and explains the spatiotemporal control of many biochemical reactions within a cell [14, 15]. The proteins within these condensates do not adopt specific conformations (i.e. they still behave like random coils) [16, 17] and thus evolve without structural restraints. Although multiple sequence alignment may work in some instances (for example, the aromatic residues in the IDRs of TDP-43 and FUS are conserved, highlighting their potential importance for LLPS [18]), most IDRs cannot be aligned, especially when there are sequence gaps between orthologs [19].

The functionally important physicochemical properties of IDPs/IDRs encoded in their primary sequence may be retained during evolution. Aromatic residue patterns [20], prion-like amino-acids [21], charged-residues blocks [22], and coiled-coil content [23] all contribute to LLPS, but these features cannot be revealed by sequence alignment. Multiple sequence alignment methods are, therefore, of limited use in identifying critical residues in IDRs. To overcome this challenge, we propose an unsupervised contrastive machine learning model trained using self-attention neuronal networks on human IDR orthologs. Our results show that the trained model “pays attention” to crucial residues or features within IDRs. We also provide online access to our model that uses primary sequences as input.

## Methods

### Training dataset preprocessing

Human protein sequences were retrieved from UniProt [24] and the corresponding orthologs were obtained from the Orthologous Matrix (OMA) database [25]. Chordate orthologs were aligned using Clustal Omega [26]. The PONDR [27] VSL2 algorithm was used to predict the IDR of the human proteins and to define the boundaries of the aligned sequences (Figure 1A). Aligned regions were defined as subgroups. N-terminal methionines were removed to assist learning (methionine is coded by the start codon in protein synthesis). After removing gaps within the aligned sequences, all sequences were padded to a length of 512 amino acids (repeating from the N-terminus; Figure 1A). The few sequences longer than 512 amino acids (56,086 out of 2,402,346, 2.3 %) were truncated from the C-terminus. The training dataset thus consisted of 28,955 ortholog subgroups from 13,476 human protein families with IDRs longer than 40 amino acids.

**Figure 1.**
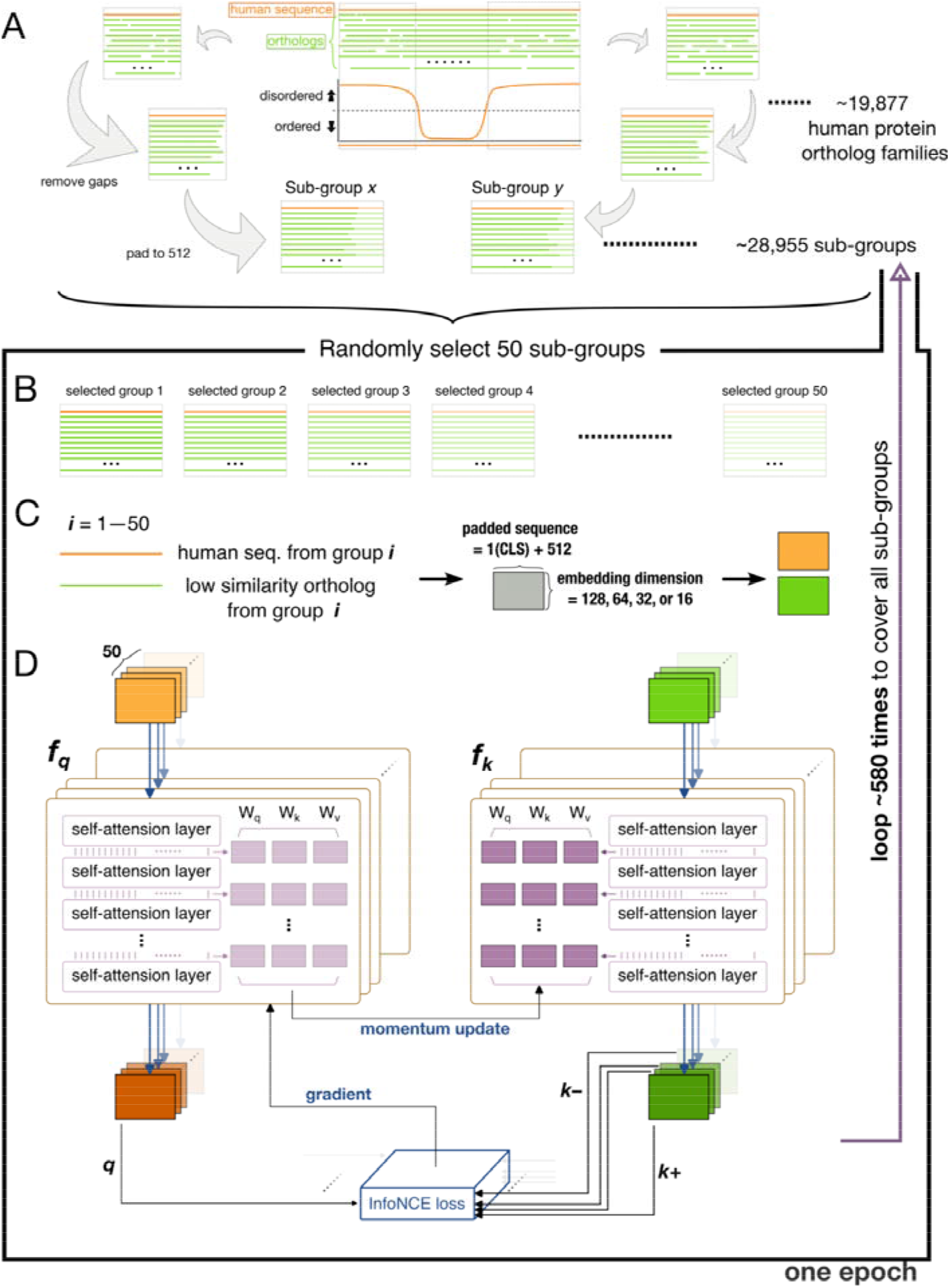
Flowchart of the training scheme. (A) Schematic representation of how the training datasets were constructed from human sequences (orange lines) and orthologs (green lines). (B) A training batch made up of 50 randomly selected subgroups. (C) Embedding of the human sequence and one of its orthologs from the same subgroup (selection probability weighted by dissimilarity) to different dimensions (as a tensor for each sequence). (D) The architecture of the training model. The steps in panels B–D were repeated 580 times to cover all subgroups in the training set, and the whole process (a training epoch) was repeated 400 times.

Each training batch consisted of fifty randomly selected subgroups (Figure 1B). The human sequence from each subgroup was paired with one of its orthologs (one of the non-human sequences in the same subgroup, Figure 1C). The selection probability was weighted by the Levenshtein distance [28] from the human sequence to favor low similarity pairings. Supplementary Figure S2 shows how different the sequences typically were in these ortholog subgroups, along with the corresponding selection probabilities. The most dissimilar sequences (high probability of being selected for training) in each ortholog group were also deposited in Open Science Framework. A classifier token (CLS) was added to the start of the selected sequences, and these were mapped to a matrix with an embedding dimension of 128 (*embed_dim;* Figure 1C).

### Training architecture

The training architecture was a self-supervised contrastive learning model, Momentum Contrast version 3 (MoCo v3) [29]. The base encoder in MoCo v3 was replaced with a classical self-attention network [30]. We used 8-head attention and tested six attention layers. Fifty human sequences from the same batch and their corresponding orthologs (the ones with the lowest similarity to each human sequence, as mentioned above) were sent to the momentum encoders (*f_q_, f_k_* respectively, following the original nomenclature [29]), and calculated in parallel (Figure 1D). The outputs from each human sequence and its ortholog were a query (*q*) and key (*k*+; the positive sample for each query). The output of the other 49 orthologs were the negative samples (*k*–). All 50 combinations of *q*, *k*+, and *k*– were formulated to minimize a contrastive loss using the adopted InfoNCE [31]:

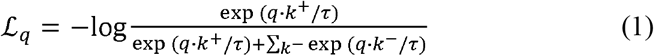

where *τ* is a temperature hyper-parameter (set to 0.02). The loss was computed in a symmetrized manner [29], i.e. the human sequences (*q*) were also sent to *f_k_*, and the orthologs (*k*) were sent to the *f_q_* with correspondent outputs for calculating the InfoNCE loss. The parameters between the attention layers of *f_q_* (light purple blocks in Figure 1D) were updated according to a gradient to minimize the cross-entropy loss (Equation (1)). The parameters in *f_k_* (dark purple blocks) were updated by the momentum encoder: (1–*m*) • query_encoder + *m* • key_encoder, with *m* set to 0.999 by default [29]. This scheme (Figure 1B–D) was repeated ~580 times to include all 28,955 subgroups in each training epoch. The training consists of 400 epochs, and the InfoNCE loss is sufficiently converged (Supplementary Figure S3). The model was built on PyTorch and the training was performed on a Nvidia Telsa P100 16G GPU.

## Results

### The trained model attributes a high attention score to experimentally confirmed critical residues

Studies have shown that the aromatic residues (phenylalanine, tyrosine, and tryptophan) in the IDRs of TDP-43 [32], FUS [33], and hnRNP-A1[34] are critical for LLPS-related functions. These residues obtain a high attention score in our model (Figure 2A). The aromatic residues (two tryptophans and ten tyrosines) in galectin-3 [35] also score highly (Figure 2B, left panel). Interestingly, although zebrafish galectin-3 differs substantially in primary sequence from human galectin-3 (Supplementary Figure S4), the aromatic residues (mostly tryptophan instead of tyrosine) also have high attention scores (Figure 2B, right panel). Note that zebrafish galectin-3 was not in the OMA ortholog database used for training (OMA number: 854142). Charged residues (purple arrows in Figure 2C) reported to be associated with condensation in NPM1 [36], FMRP [37], and Caprin1 [38] also obtain high attention scores (Figure 2C). Our model also assigns high attention scores to the methionines in Pbp-1 (labeled in Figure 2D; Pbp-1 is the yeast ortholog of human Ataxin-2), which have been shown to be critical for redox-sensitive regulation [39]. Altogether, these results indicate that the trained model correctly identifies known key IDR residues.

**Figure 2.**
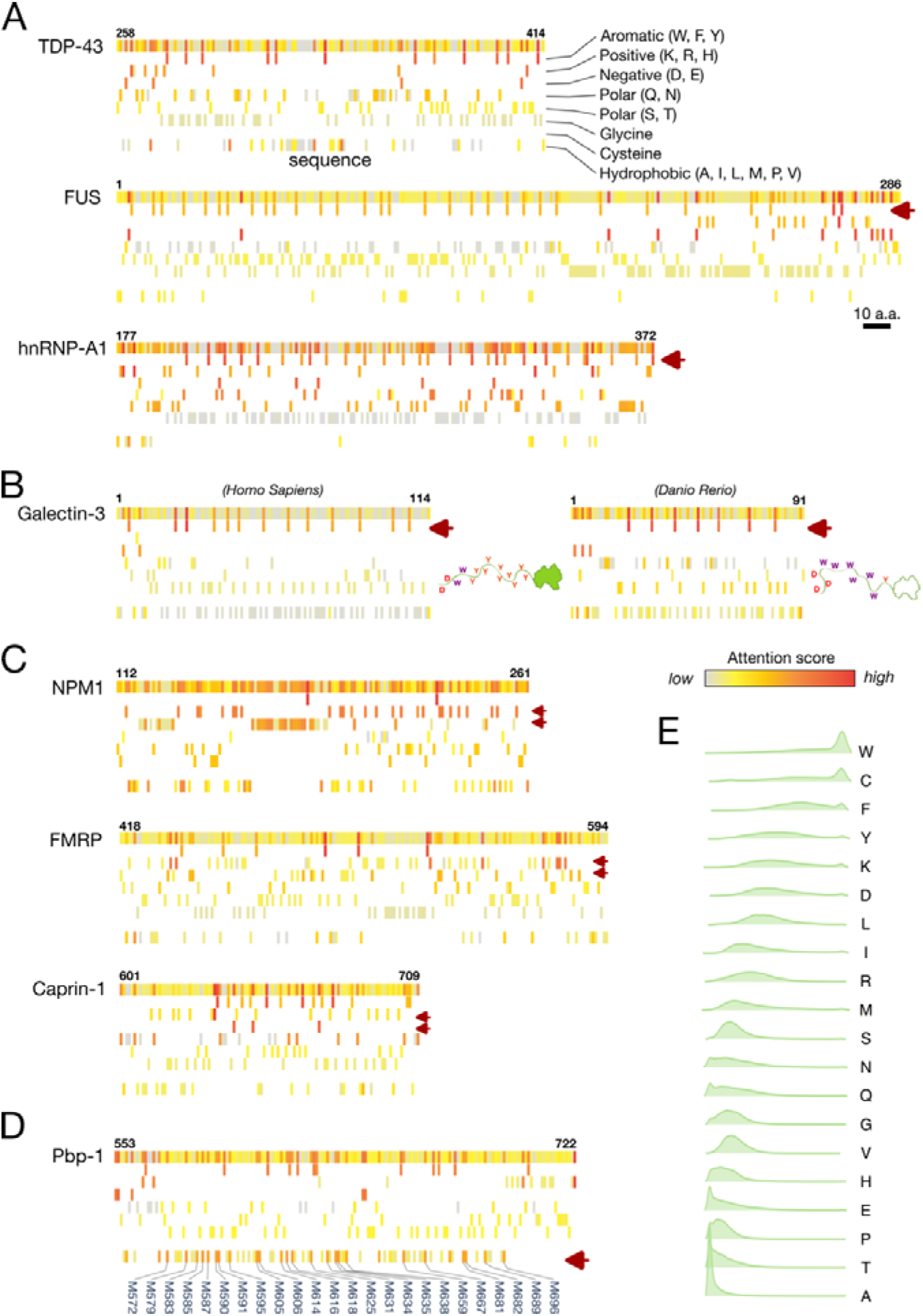
Results of the trained model for reference proteins and attention score distributions for individual amino acids. (A–D) Sequences and attention scores for the intrinsically disordered regions of (A) the RNA-binding proteins TDP-43, FUS, and hnRNP-A1, (B) human and zebrafish galectin-3, (C) NPMA, FMRP, and Caprin-1, and (D) Pbp-1. The attention scores appear as heatmaps from high (red) to low (grey) in the top row of each protein along with residue numbers. Amino acids with different physical properties are shown on separate rows as indicated in panel (A). Purple arrows indicate amino acids of known functional importance. (E) Half-violin plots of the distribution of attention scores in human IDRs for each amino acid, sorted by median value from high (tryptophan, W) to low (alanine, A).

### Most amino acids have broadly distributed attention scores except tryptophan and cysteine, whose presence in IDRs hints at potential importance

Figure 2E compares the attention score distributions of the amino acids in human IDRs. The differences are striking, but the attention scores are not correlated with other physical properties, such as disorder/order propensity [40, 41], prion-likeness [42], or prevalence in human IDRs (Supplementary Figure S5). The attention scores of alanine are always low. Although alanine promotes α-helix formation, which is known to contribute to IDR functions such as LLPS [43–45], our model ignores these residues. This is probably because α-helices are also promoted by other amino-acid types, e.g. leucine or methionine [46, 47], in different combinations not involving alanine. Also, since the training process did not include structure information, structure-related sequence motifs were ignored. At the other end of the distribution, tryptophan and cysteine systematically obtain high attention scores. These structure-promoting amino acids rarely appear in unstructured regions [40, 41, 48]; therefore, their appearance in IDRs hints at their potential importance. Although little is known about the role of cysteine in IDRs, its involvement in tuning structural flexibility and stability has been recently discussed [49]. Tryptophan, in contrast, is well-known to act as LLPS-driving “stickers” in IDRs [32, 50, 51], and bioinformatic analysis shows that they may have evolved in the IDRs of specific proteins to assist LLPS [18].

Finally, the fact that most amino acids, including those highlighted in Figure 2A–D, have broad attention score distributions (Figure 2E), excludes the possibility that our model is biased toward particular amino-acid types rather than sequence content as a whole. Moreover, in the machine learning procedure, the protein sequences were embedded into higher dimension matrices (as sequences of digits; Figure 1C), and amino-acid type information was lost when the matrices were transformed into tensors along with the self-attention layers (Figure 1D). These results support the predictive ability of the trained model.

## Discussion

Genetic information, in the form of a linear combination of nucleic or amino acids, becomes more diverse over time. Comparing levels of diversity between different species reveals how closely related they are. In terms of amino acids, multiple sequence alignment not only highlights phylogenetic relationships between proteins but also facilitates homology modeling for structure prediction [52–54]. Machine learning approaches have recently been used to incorporate information from evolution to train structure prediction models [55, 56], and the highly accurate predictions from AlphaFold [57] and RoseTTAFold [58] have revolutionized structural biology. In contrast, the structural conformations of IDRs do not have a one-to-one correspondence with the primary sequence, and multiple sequence alignment often fails [18, 59]. These limitations make IDR structural ensembles challenging to predict. A few attempts have been reported, such as using generative autoencoders to learn from short molecular dynamics simulations [50]. The potential and challenges of machine learning in IDR ensemble prediction are also discussed [59].

Sequence pattern prediction faces similar challenges, including the lack of a sufficient stock of “ground-truth” training data, such as image databases or the Protein Data Bank. However, unsupervised learning architectures have been developed to train models without labeled datasets [60], and this type of approach is especially well-suited for IDRs. For instance, Saar et al. used a language-model-based classifier to predict whether IDRs undergo LLPS [61]. Moses and coworkers pioneeringly applied unsupervised contrastive learning, using protein orthologs as augmentation [62], to train their model to identify IDR characteristics [63]. Although we also used ortholog sequences as training data, our approach differs in many aspects. Instead of convolution neural networks, we used self-attention networks to capture the distal features in the entire protein sequence. Additionally, we trained our model using the latest contrastive learning architecture (MoCo v3), which greatly reduces memory usage for larger batches and enhances efficiency. In contrast to other masked language models [64–66], our approach is the first, to the best of our knowledge, to combine contrastive learning and self-attention in extracting features using natural language processing for protein sequence analysis. Our trained model directly “pays attention” to potentially critical residues in the entire sequence instead of mapping the primary sequence to learned motifs [63]. In other words, our model identifies overall features in an IDR sequence, for example, a predominance of aromatic residues or blocks of charged residues (Figure. 2). Moreover, our model provides intuitive results that point out potentially important residues for researchers to target for example in mutagenesis or truncation experiments.

## Conclusion

Although the model could be improved by training on larger datasets (e.g., including more orthologs other than human’s) or with larger batch sizes (requiring a supercomputer), these results show that self-supervised contrastive learning with self-attention networks can be used to identify key residues in IDRs, something that cannot be achieved by conventional multiple sequence alignment. The model, IFF – for *I*DP *F*eature *F*inder, can be accessed online using a primary sequence as the only input. We expect our model to be useful in various research fields, notably cell biology, to efficiently identify critical residues in proteins with IDRs, such as those that undergo LLPS.

## Acknowledgments

The authors thank the IT Service Center at NYCU for GPU access. This work was supported by the National Science and Technology Council of Taiwan (110-2113-M-A49A-504-MY3). The authors are also grateful to Prof. Wen-Shyong Tzou (National Taiwan Ocean University) for his help.

