## Supporting Information for "Identifying key residues in intrinsically disordered regions of proteins using machine learning"

for

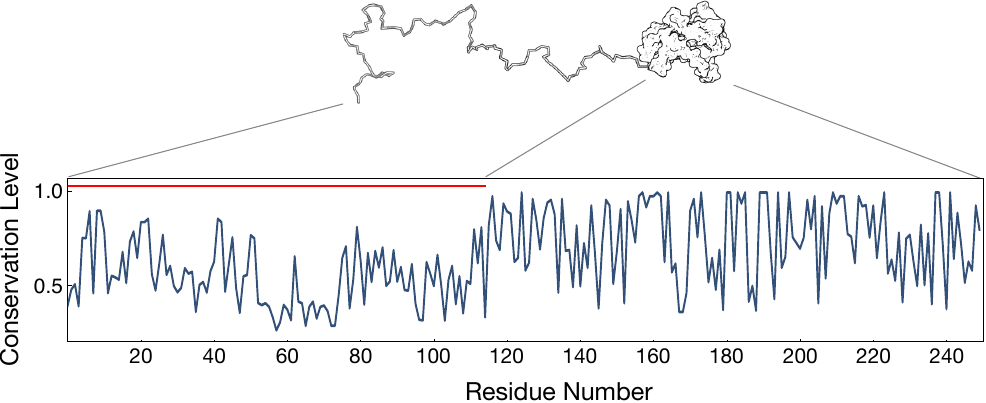

**Figure S1**. Sequence conservation in the folded and intrinsically disordered regions of galectin-3 among chordates in the OMA database (access number: 854142; 92 sequences analyzed). Sequence conservation is expressed in terms of the Shannon sequence entropy [1] of each residue. Conservation levels are lower in the disordered region (highlighted by a red horizontal line).

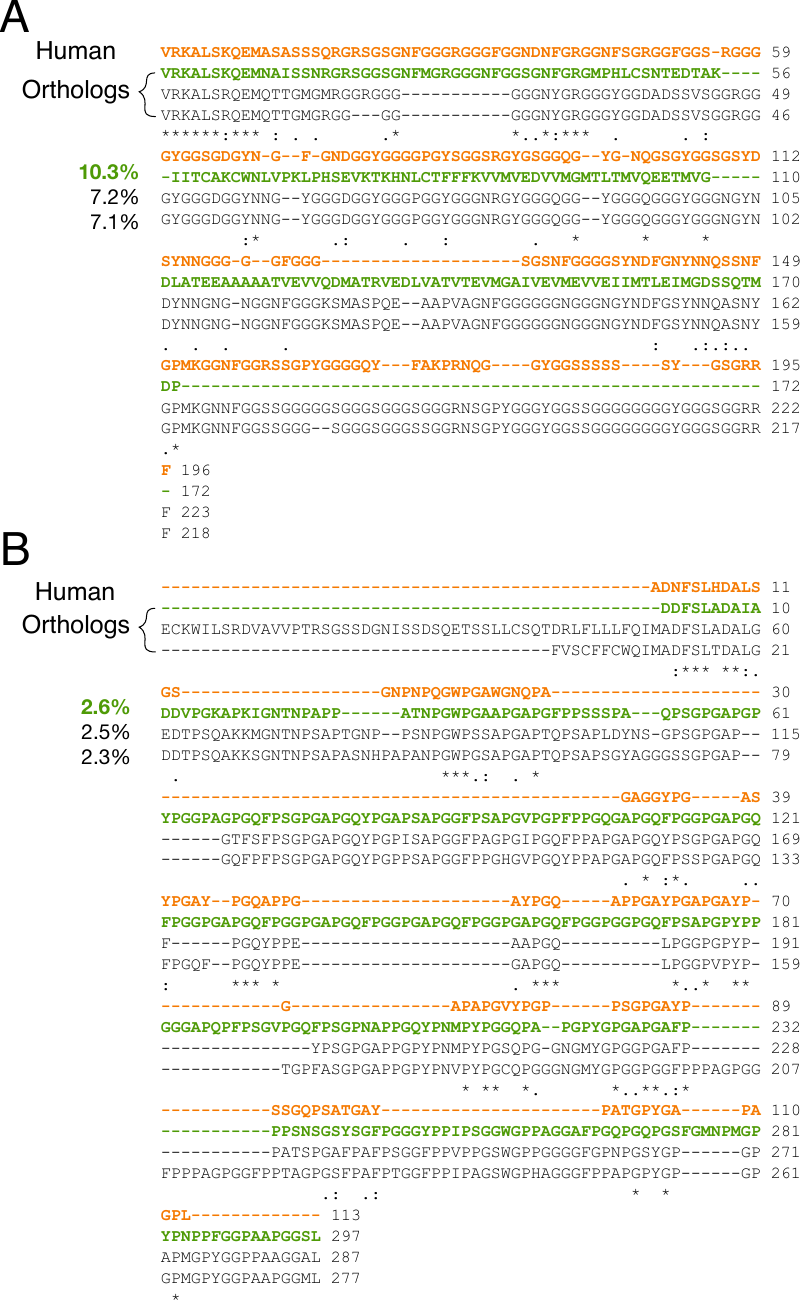

**Figure S2.** Comparison of ortholog sequences in a training set. (A) hnRNPA1 and (B) galectin-3. The human sequence is shown in orange along with three orthologs with low similarity (high Levenshtein distance). The most dissimilar ortholog sequence (the one with the highest selection probability, indicated on the left) is shown in green.

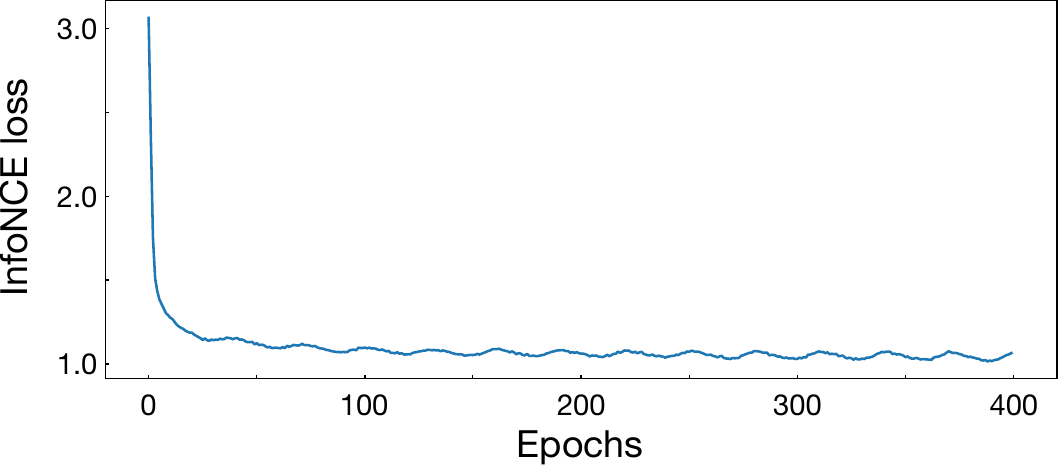

**Figure S3.** InfoNCE loss as a function of epoch iterations for the training process illustrated in Figure 1 in the main text.

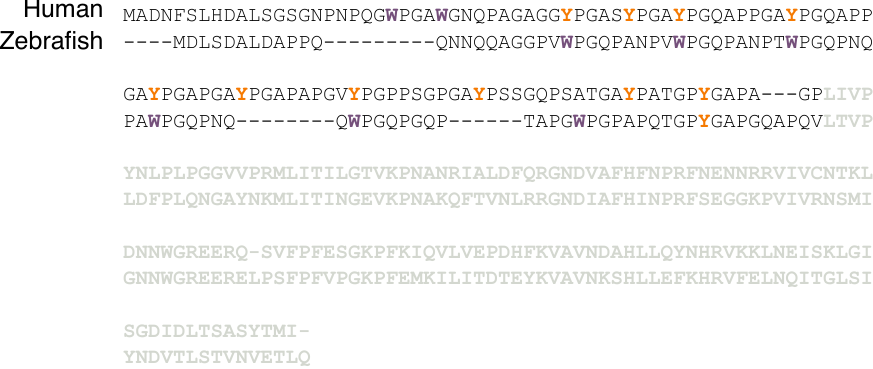

**Figure S4.** Alignment of human and zebrafish galectin-3 sequences. The UniProt entries are P17931 (*Homo Sapiens*) and Q6TGN4 (*Danio Rerio*). Amino acids in the structured domain are shown in grey. The aromatic residues in the intrinsically disordered regions are colored purple for tryptophan (W) and orange for tyrosine (Y).

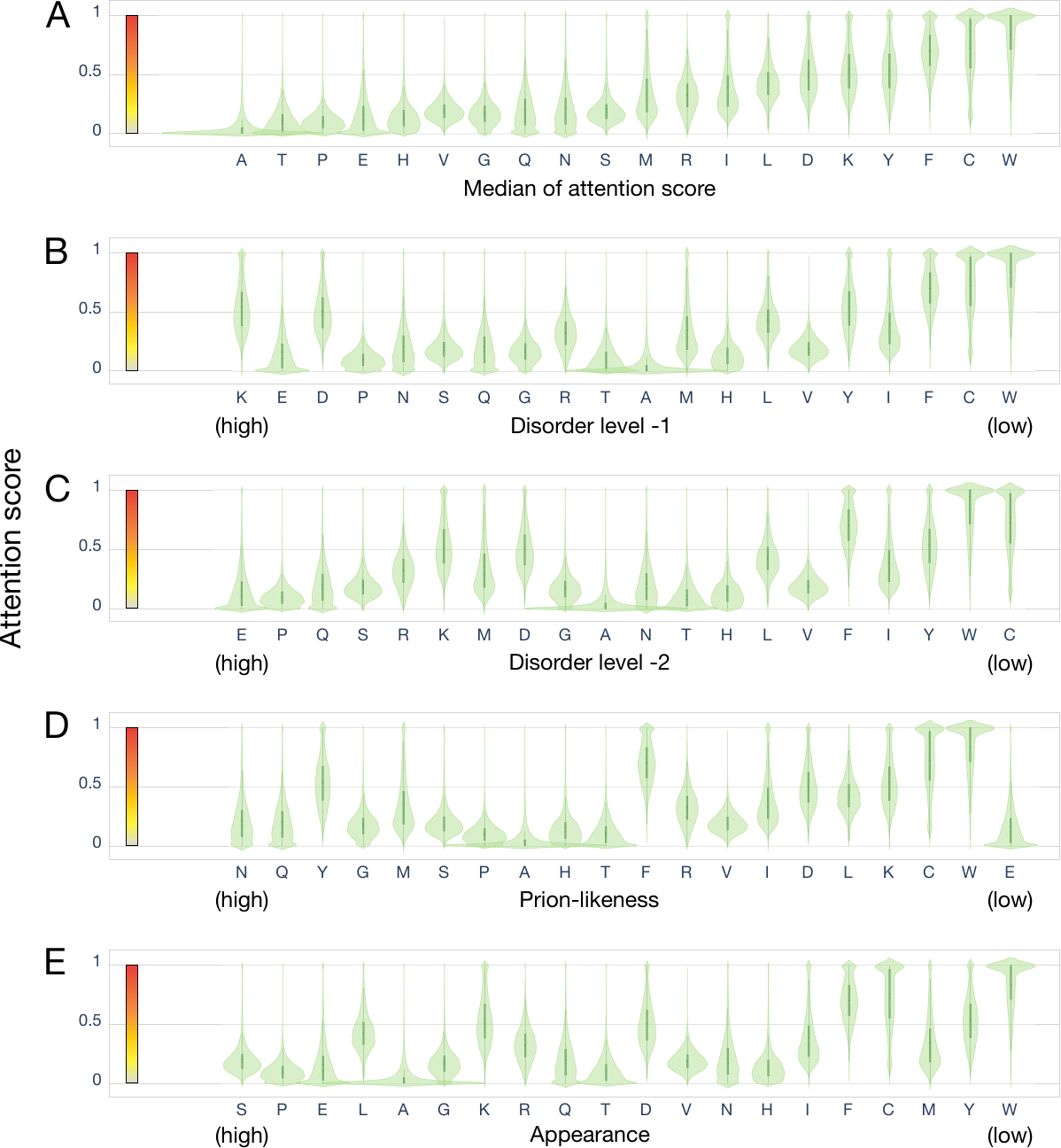

**Figure S5.** Violin plots of the distributions of attention scores for each amino acid, with the amino acids sorted by different physical properties along the horizontal axis. (A) Sorted by median attention score as in Figure 2E in the main text. (B, C) Sorted by preference in disordered regions, according to (B) Vihinen et al. [2] or (C) Radivojac et al. [3]. (D) Sorted by prion-likeness [4]. (E) Sorted by prevalence in the human IDRs used for training. (S: ~402k times, P: ~334k, E: ~330k, L: ~280k, A: ~269k, G: ~261k, K: ~237k, R: ~235k, Q: ~206k, T: ~194k, D: ~172k, V: ~148k, N: ~113k, H: ~88.2k, I: ~87.9k, F: 72.4k, C: 65.2k, M: 64.3k, Y: 52.3k, W: 21.5k).
